# Rapid classification vs. refined learning about harm intent: Roles of serotonin and racial bias

**DOI:** 10.1101/2023.05.20.541280

**Authors:** Michael Moutoussis, Joe Barnby, Anais Durand, Megan Croal, Laura Dilley, Robb B. Rutledge, Liam Mason

## Abstract

Attributing motives to others is a crucial aspect of mentalizing, can be biased by prejudice, and also is affected by common psychiatric disorders. It is therefore important to understand in depth the mechanisms underpinning it. Toward improving models of mentalizing motives, we hypothesized that people quickly infer whether other’s motives are likely beneficial or detrimental, then refine their judgment (‘Classify-refine’). To test this, we used a modified Dictator game, a game theoretic task, where participants judged the likelihood of intent to harm vs. self-interest in economic decisions. Toward testing the role of serotonin in judgments of intent to harm, we delivered the task in a week-long, placebo vs. Citalopram study.

Computational model comparison provided clear evidence for the superiority of Classify-refine models over traditional ones, strongly supporting the central hypothesis. Further, while Citalopram helped refine attributions about motives through learning, it did not induce more positive initial inferences about others’ motives. Finally, model comparison indicated a minimal role for racial bias within economic decisions for the large majority of our sample. Overall, these results support a proposal that classify-refine social cognition is adaptive, although relevant mechanisms of Serotonergic antidepressant action will need to be studied over longer time spans.

## Introduction

Relationships with others play a key role in well-being and social harmony; to wit, mental health suffers when important relationships deteriorate. Clinically, perceiving others as harmful is central in conditions ranging from PTSD (assaults) to social anxiety (humiliation) to paranoia (conspiracy). It is thus important to understand in depth the ways in which perceptions of others, especially regarding under-privileged groups (Kaiser Trujillo et al., 2022; Singh et al., 2022), may interact with mental well-being.

The ’Bayesian Brain Hypothesis’ holds that confidence in beliefs, even those characterizing paranoia or PTSD, is updated by the brain (approximately) employing Bayes’ rule. How strong one ‘belief’ is depends on conviction in related ones, so beliefs become organized into interconnected hierarchies, ranging from simple predictions regarding sensory data to abstract expectations about self and others encoding high-level features from the environment. Belief-based models account well for data related to harm attribution, over and above associative learning models (Barnby et al., 2020, 2022), but much remains unclear. For example, understanding the neuro-computational basis of polarized attributions about others is at an early stage (Brown et al., 2022; Story et al., 2023).

The psychological literature has traditionally casted polarized thinking as maladaptive, contributing to us-them biases. However, taking inspiration from recent modeling work (Story et al., 2023), here we ask whether polarized attributions may originate in useful adult cognition.

We propose a “classify-refine” hypothesis, whereby people first (1) attempt to rapidly *classify* others’ attributes as ‘beneficial vs. detrimental’ to themselves, and subsequently (2) to *refine* beliefs about others through learning. This may be highly adaptive: quickly telling friend from foe, computationally easy, then allowing one to refine their beliefs and go beyond black-and-white thinking. We aimed to test this classify-refine hypothesis, and improve upon limitations of existing work (Bone et al., 2021; Hopkins et al., 2021) through the models overviewed in Fig. 2. We compared ‘classify-refine’ models, that used only one beneficial and one detrimental state along each dimension of attribution, which classic learning models employing a fine-grained range along each.

Social experiments based on game theory have been useful in modeling cognitive processes of inter-personal inference (Barnby et al., 2020, 2022; Greenburgh et al., 2019; Raihani & Bell, 2017). We extend this work towards a more ecologically valid task for probing brain mechanisms behind attributions, based on the repeated Dictator Task (Barnby et al., 2020, 2022). Here, a “dictator” (an on-screen partner of the participant) decides how to split a sum of money between themselves and a “receiver” (the participant), who must accept the split. The motivation of the "dictator" is undisclosed, but receivers rate, for each economic exchange, the extent to which decisions may have been motivated by harmful intent vs. the extent to which they were motivated by self-interest.

Work with this attribution task has not as yet studied some important social variables. One variable thought to influence beliefs in real life is race or ethnicity. For example, the implicit association task has consistently demonstrated implicit race-related biases, which frequently but not invariably influence behavior (Maina et al., 2018). Toward querying a possible role of race in attributions of intent, here we implemented the task using photos of either black or white individuals to portray “dictators”.

Depression is associated with a ‘hostile interpretation bias’, i.e., increased propensity to interpret ambiguous behaviors as hostile (H. L. Smith et al., 2016). Anecdotally, we have encountered clinical cases of amelioration of racially hostile attitudes in patients suffering from Alzheimer’s disease and depression during treatment with antidepressants. Attribution of harmful intent is implicated in anti-black racism (Stjohn & Healdmoore, 1995). Strikingly, negative emotionality has been reported to mediate the impact of adverse events on bias against out-groups (She et al., 2022). The commonest antidepressants in use are the selective serotonin reuptake inhibitors (SSRIs), such as Citalopram. In the presence of Citalopram therapy, relationship factors are associated with improvements in depressive symptoms (Joseph et al., 2011). Hence, antidepressants may reduce attributions of harm intent, but whether they do so is unknown. We hypothesized that SSRIs may reduce attributions of harm intent, and to test this, we administered the repeated dictator task before and after subchronic Citalopram treatment.

In summary, the present study aimed to shed light on the neurocognitive basis for how individuals gauge harm vs. self-interest using Bayesian modeling of a repeated Dictator task that we call the Sharing Game. This generated data for testing whether in interpersonal situations, individuals initially quickly classify others’ attributes as beneficial vs. detrimental to themselves, and subsequently refine and update their beliefs (“classify-refine hypothesis”). We employed two experimental manipulations to test two pre-registered predictions. First, we manipulated serotonin levels by randomizing participants to receive Citalopram vs. placebo. We predicted that Citalopram would result in attributing more beneficent motives. Second, we evaluated whether race plays a role in evaluations of harm vs. self-interest by using photos of white and black ’dictators’. We predicted that more negative attributions would be made for out-group others (Moutoussis et al., 2022b) i.e., those identifying as white would make more negative attributions for non-whites than whites, while non-whites would evaluate non-whites more positively than whites.

## Materials and Methods Sample

Healthy UK residents were recruited from a University College London (UCL) subject pool. They gave informed consent to participate in a week-long Citalopram 20mg vs. placebo study, approved by the UCL Ethics Committee, ID 19601/001. Participants had no history of psychiatric or neurological disorder and agreed not to become intoxicated by drugs or alcohol during the study. Seventy-four participants were enrolled (44/74 self-identified Female, 29/74 Male, 0/74 other, 1 missing). 42 participants were randomized to Citalopram. The commonest ethnicities were white and Chinese. The sample was highly educated, young adult (median age = 25), and of low income (Supplemental Fig. S2).

### Task

In our Sharing Game task (Fig. 1A), a development of the repeated dictator task (Barnby et al., 2020, 2022), we increased task length by one block, we collected more data per trial, and tested a new set of computational models. In so doing we aimed to increase the stability and sensitivity of tasks, toward improving the assessment of individual differences, including drug effects.

**Fig. 1.**
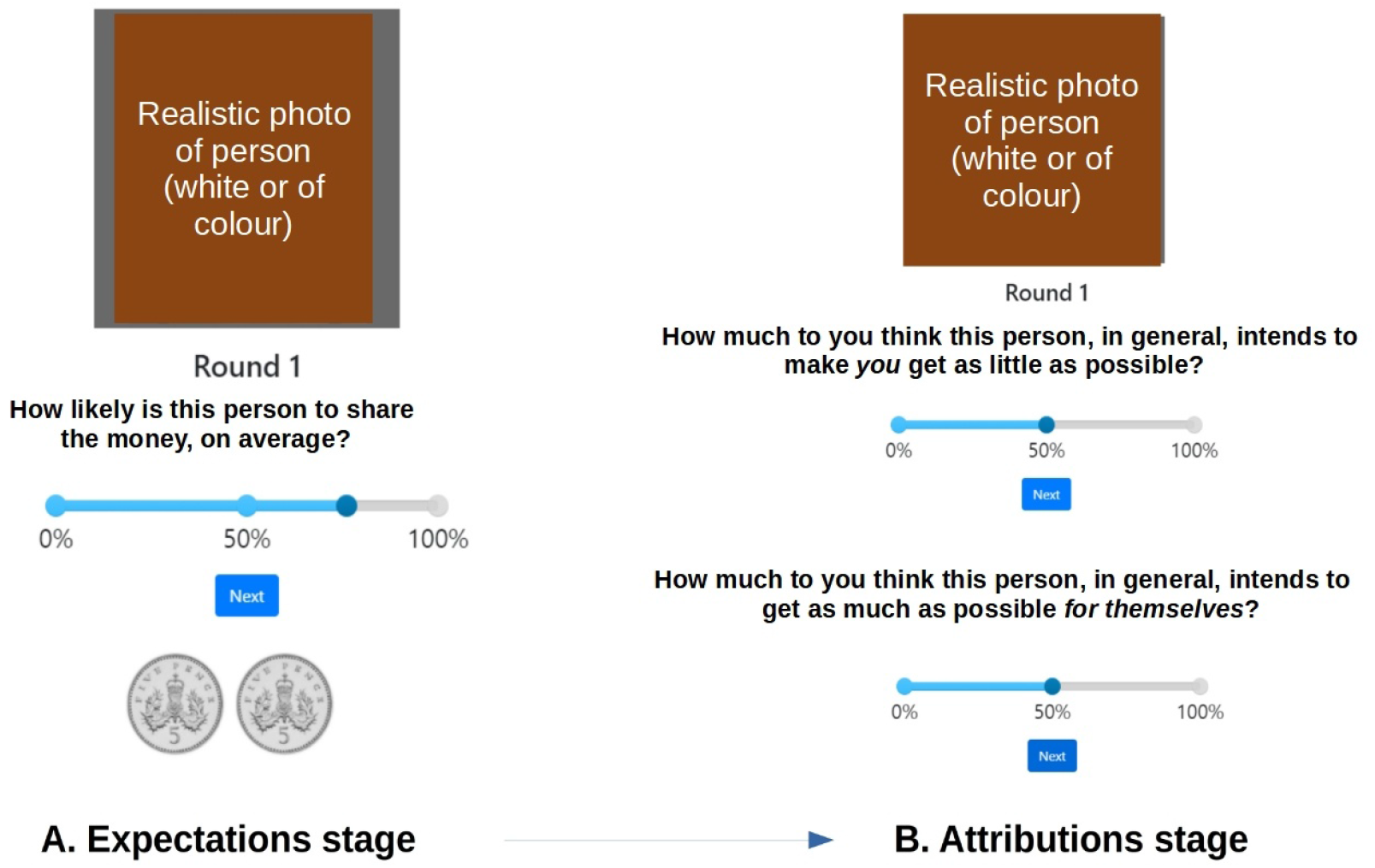
One trial of the revised iterated dictator task, a.k.a. ’Sharing game’. **A.** Expectation stage. Participants were asked to estimate the probability that this Dictator would split two coins fairly. **B.** Having observed the split (not shown here), participants had to infer the likelihood of Self-interest and Harm-intent motivating the Dictator. Question order was randomized across trials. The expectation stage preceded attributions within a trial. Computationally, it is best thought of as the result of belief updates formed on the basis of the observations made so far, especially in the trial before. Unlike many economic games, we displayed ecologically valid Dictator images, and asked whether they elicited motivational attributions more beneficial or detrimental to the participant.

Participants thus saw four Dictators, who made either fair (5:5) or unfair (10:0) splits of 10 pence of fictive money between themselves and the participant (2 Dictators 80% fair, 2 Dictators 20 % fair). Participants reported their expectations about the fairness level of the Dictator, then made attributions of motivation along two salient dimensions: Harm-intent (HI) and Self-interest (SI). (Fig. 1). The dictators were portrayed by photos of women of varying age (young adult to middle-aged) and ethnicity (black vs white), purchased under license for public use from www.shutterstock.com .

73 participants completed the Sharing Game at least once; sixty-six completed it again one week later. They faced each dictator just once, for 12 consecutive trials. A post-experiment survey probed awareness of the drug and its subjective effects. Of the 42 Citalopram participants, 25 experienced subjective drug effects (all minor) and correctly guessed that they took SSRI.

### Modeling

All models had a simple hidden Markov process (HMM) as a central ‘learning core’, implemented as a one-level, 12-trial HMM. Each model had three ’reporting’ processes, generating fairness expectation, Harm-intent attribution and Self-interest attribution reports. All models were derived from a published, successful Bayesian model (Barnby et al., 2022). Here we used the active-inference framework, which naturally accommodates the process of fast classification, accompanied by slower belief refinement (R. Smith et al., 2022). We now describe the key features of the models (Fig. 2). The parameters are explained in Table 1., and their place in the different models which were compared in Fig. 4. Detailed explanations and equations follow in the Supplement, where Fig S1 exemplifies the workings of the classify-refine model.

**Fig. 2.**
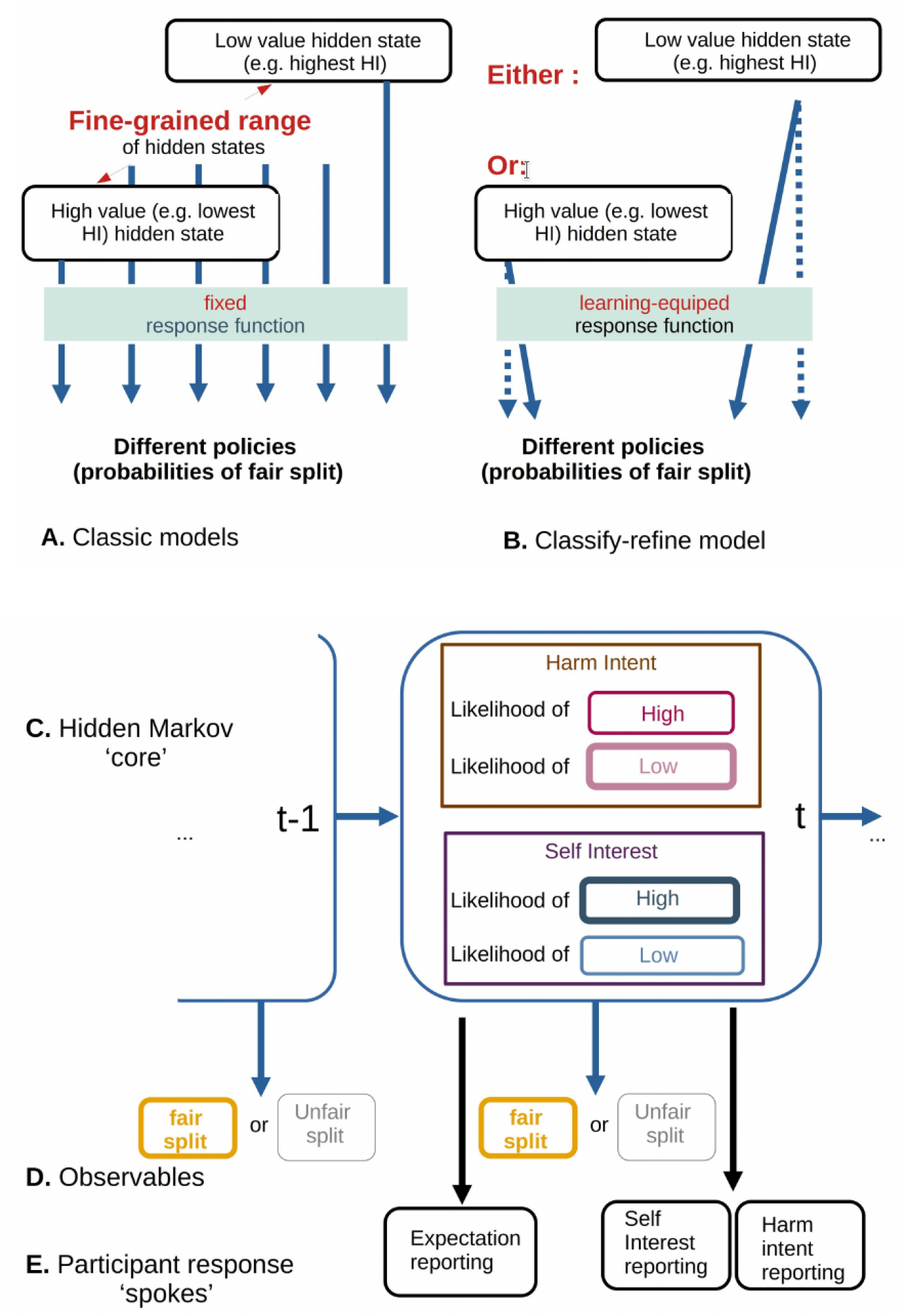
Essentials of Classic vs. Classify-refine models **A.** In classic models, each of a fine-grained-range of states maps to a specific policy. E.g., the worst possible Harm Intent always maps to an ’unfair splitting’ polity. Participants have to infer which of these many states obtains. **B.** In the new model, classification into coarse-grained values occurs, i.e. only detrimental and beneficial, but what these mean for each partner is refined through learning. **C.** Beliefs about states form the ’core’ of the generative model, updated at each trial. **D.** Returns seen at each trial. **E.** Participants rated their expectation of fair or unfair split before observing the return, and afterwards reported their (updated) attributions. Although they consider binary states, they have uncertainty about which obtains, so they still report a graded score about how likely a particular attribution is.

**Fig. 3.**
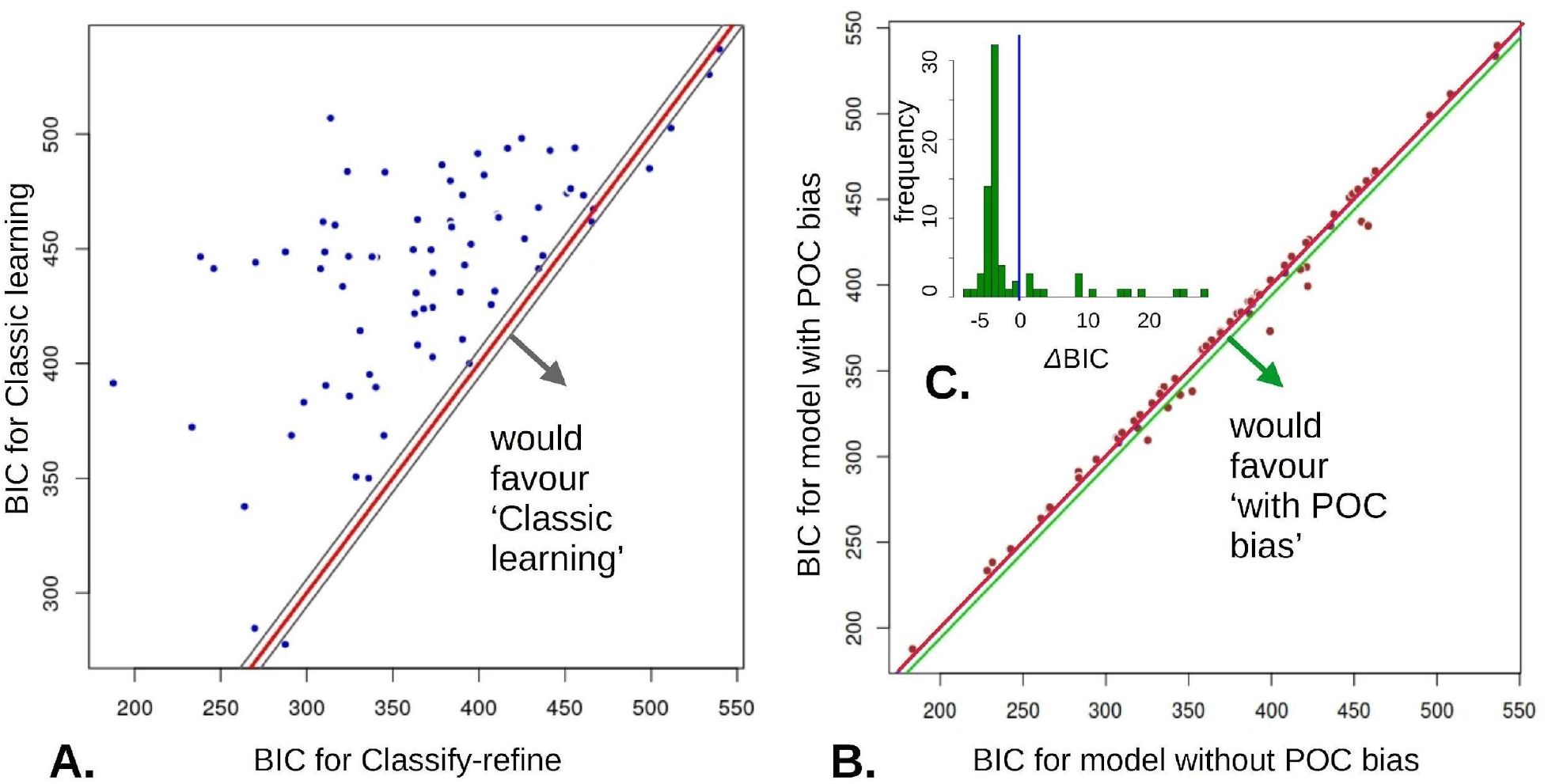
Key model comparison results. Red lines = BIC equality. **A.** Classify-refine models clearly outperform classic learning. For 64/73 participants, the BIC difference is 6 or more (i.e. to the left of gray, diff = +6 gray line; conventionally at least modest evidence. Right gray line is diff = -6). Blue line is identity. **B, C.**: Race / ethnicity bias. **B.** The model without person-of-color bias gave a more parsimonious fit for 63 out of 73 participants (dots above the equal BIC line). For 10/73 participants, the model including a POC bias parameter gave an advantage of at least 6 BIC points (‘modest evidence’) **C.** (inset) Histogram of the difference, showing a main peak for the simpler no-bias model and an upper tail for the more complex model.

**Fig. 4.**
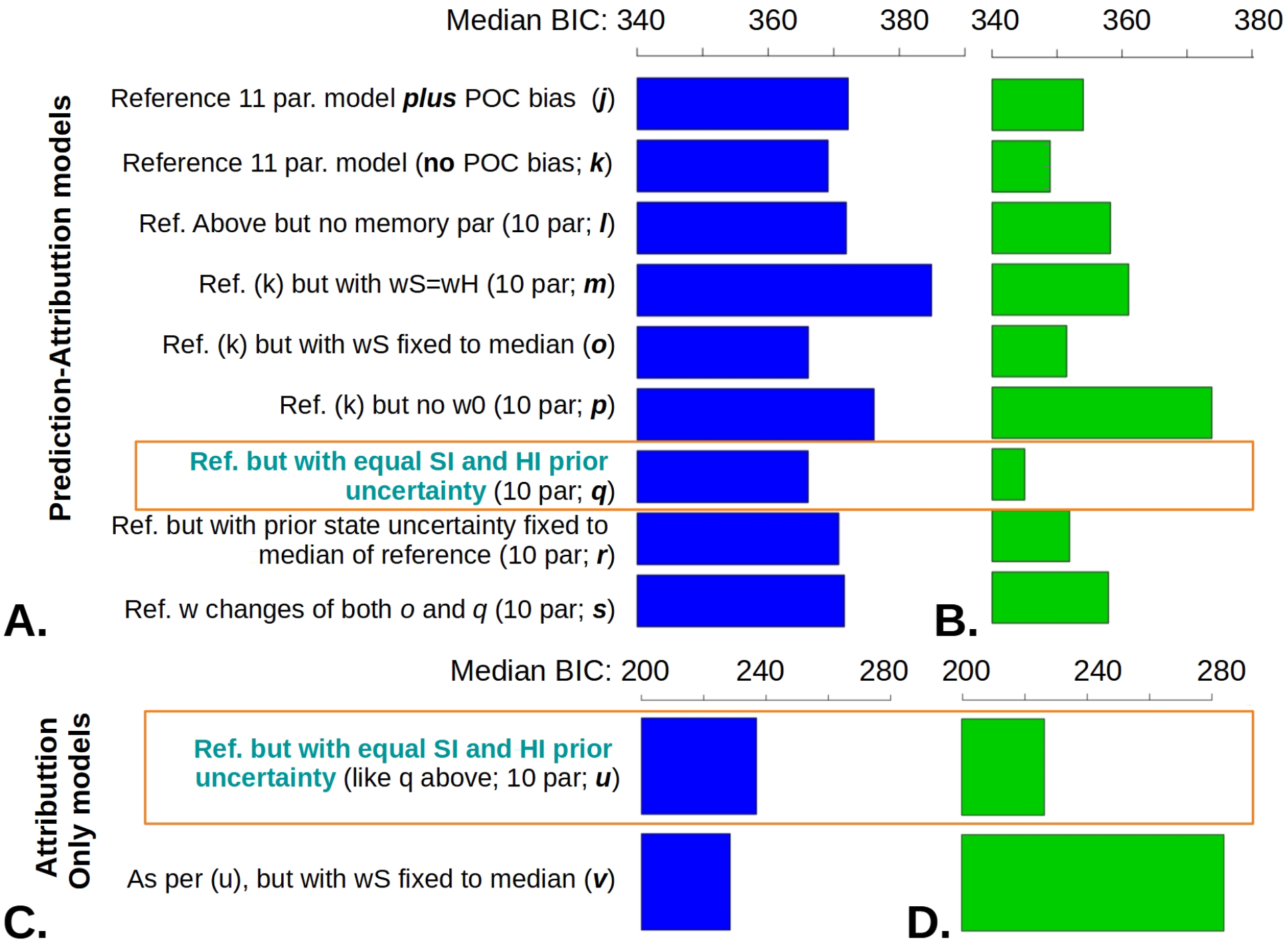
Median Model Fit measures (BIC) for Classify-refine models. **A.** The best model at both baseline and follow-up (**B.**) had two parameters fewer than the full model - POC_bias and ratio of prior certainties for HI vs. SI . BIC reduced with testing wave, mixed-effects analysis BIC `∼` wave + ( `∼`1 | participant) p=0.023. Note the same scales. **C., D.** : Overall, the same model was the best when fitted only to the attribution, but not the expectation data. This was a much less stable model (cf. log-likelihoods in Fig. 3), which probably accounts for why it was slightly worse than its main competitor shown here at baseline (C.), but considerably better at follow-up (**D.**).

**Table 1.**
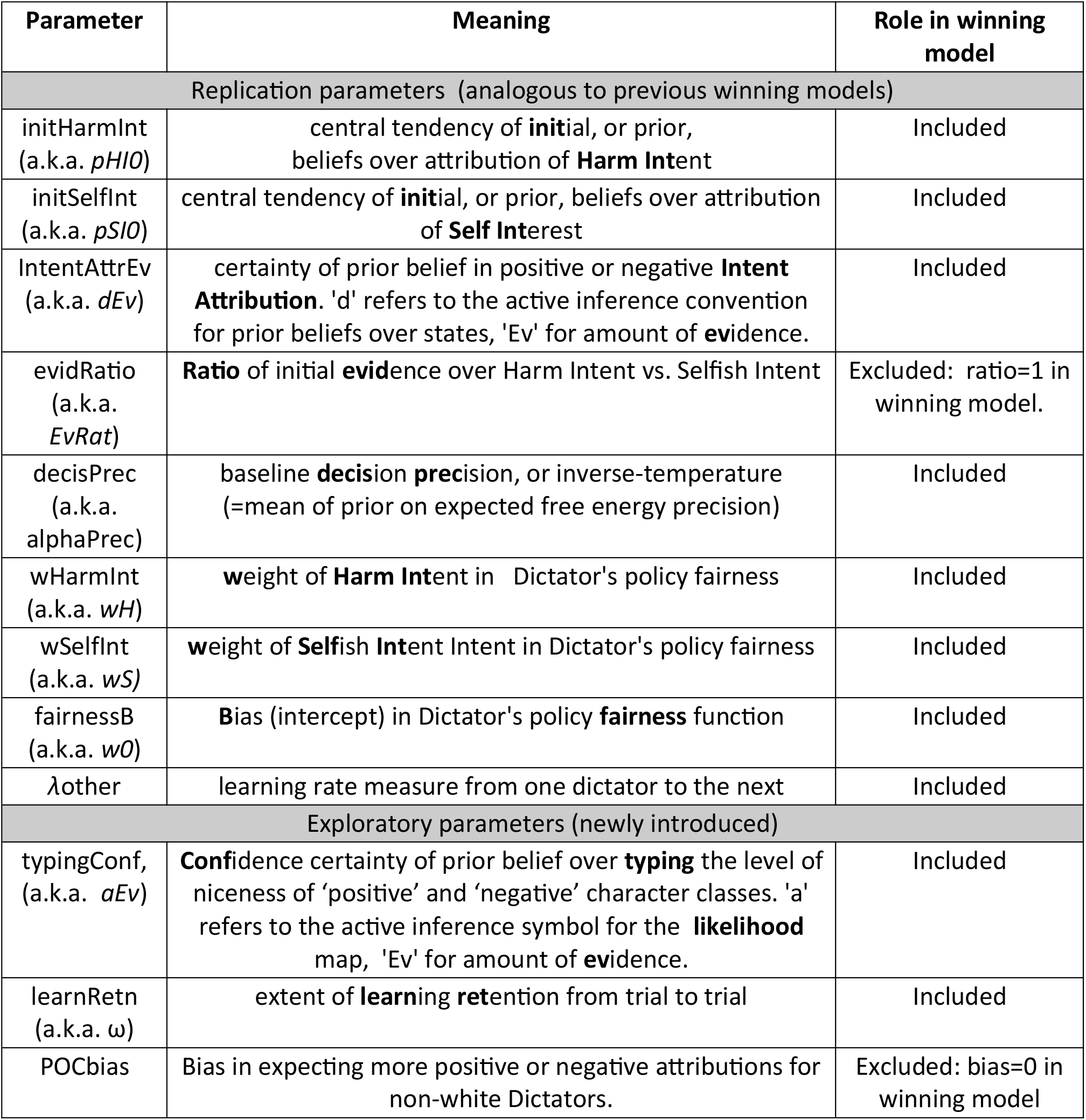
Definition and roles played by model parameters

Agents inferred the type of others over an array of Self-Interest x Harm-intent states, utilizing independently parametrized priors over each dimension (Table S1). Each ’type’ (SI, HI) determined a probability of fair-split through a logistic function (Supplement Eq. S1). Observed splits allowed agents to invert this model and update their beliefs. Beliefs held at the learning ’core’ produced ’fairness predictions’, ’HI reports’ and ’SI reports’ via three simple active- inference modules.

In Classic models, HI and SI grids had 6 bins each, with fixed values, covering the possible range; but for the classify-refine models, the central core only had two states along each attribution dimension (Fig. 2B), giving four combinations. These, however, were not of fixed value, but could be adjusted via learning. Initially, each was taken to include low (5%) and high (95%) values of each attribute, so as to represent the psychological meaning of ’beneficial’ and ’detrimental’ and span the range of each attribute. Crucially, uncertainty over the location of the bins was parametrized by a *typing confidence* parameter *aEv* (Table S1). Beliefs about location of the ’beneficial’ and ’detrimental’ states were refined by crediting the evidence for ’fair’ or ’unfair’ split at each trial in proportion to the (posterior) belief that said state underlied the trial (Dorfman et al., 2019).

Ethnicity did not directly affect learning, but modulated the beliefs fed from the learning to the reporting processes. That is, the core HMM learnt without taking race into account, but its output to the response modules was subject to a bias parameter. The magnitude and direction of this bias was fitted individually, allowing modeling of the bias in any direction - including being positively biased about an out-group.

#### Model fitting

We fitted participants’ expectations about how the Dictator would split the sum, and their attributions of the likely self-interest and harm-intending motives of the Dictator, using models parameterized as per Table 1. Maximum-a-posteriori (MAP) fitting with weakly informative priors defined over native parameter space was used (Moutoussis et al., 2018). The sum-log-likelihood at the MAP estimate was then used to calculate Bayesian Information Criterion (BIC; corrected for small samples) and Akaike IC (AIC) for each participant. In cases where a parameter was fitted across the whole participant sample, we modified the complexity penalty associated with that parameter to be consistent with BIC/2 being an approximation to log model evidence. Gradient-descent methods (including matlab *fmincon* and SPM *spm_dcm_mdp*; (matlab, 2019; SPM development team, 2022) encountered problems with local minima, hence we used adaptive grid-search optimisation with multiple initial conditions.

#### Regression analyses

Model fitting mostly resulted in approximately normal parameter distributions in transformed space, where regression analyses were performed. Sometimes, however, outliers occurred. We therefore first used robust regression *rlm* in R (R Core Team, 2020), and we report hypothesis tests based on the t-value of the coefficient in question, e.g. rlm( pHI0_follow-up ∼ pHI0_baseline + drug_group + gender + subjective_socioecon_status ). We then performed the equivalent OLS regression using *lm* and identified points with Cook’s distance > 1 and/or standardized residual distance > 3 from the theoretically predicted on Q-Q plots. We excluded these suspect datapoints from further analyses. After these few outliers were excluded, we performed longitudinal analyses using linear mixed effects models.

## Results

### Refined learning accompanies fast classification

In our baseline sample, the Classify-refine model was superior to the classical learning model (Fig. 3A. median BIC 372.2050 vs. 446.69, Wilcoxon rank sum p < 1E-5), strongly supporting our hypothesis that individuals rapidly classify others’ attributes as beneficial vs. detrimental to themselves, followed by refining their attributions. People refined their attributions more slowly if they had a higher typing confidence parameter, *aEv*. This quantified the initial amount of evidence underpinning their beliefs that each type of partner would follow a specific policy (a-matrix; see Table 1 and Methods). Further model comparisons performed on the baseline data indicated the necessity of including learning from one dictator to the next (*λ*_other_ in Table 1) and imperfect memory for retaining learning from trial to trial (*ω*). We then fixed one parameter at a time, in order to discover more parsimonious models, or replicate our previous successful models as per pre-registration. The necessity for all parameters replicated, except the need for separate uncertainties over prior Harm and Selfishness intent (analogous to uSI0 = uPI0 in our previous model, and replicating novel work (Barnby et al., 2023)). The only remarkable correlation between parameters was between *initHarmInt* and *initSelfInt* (baseline r=0.438, p uncorr. = 0.000170, follow-up r=0.428, p uncorr. = 0.000372, Fig. S8).

### Coalitional factors: lower subjective status may attenuate attributions of harm intent

To examine the pre-registered hypothesis that race stereotypes would modulate attributions, we examined the effect of a parameter that shifted the effective attributions as they were communicated from the ’core belief’ module to the ’Dictator response function’ (Fig. 2A to B), based on the apparent ethnicity of the Dictator.

Happily, we found strong evidence against our hypothesis that including this bias parameter regarding people of color (POC) would improve model fit for most participants. The model including apparent ethnicity had a median BIC of 372.2, vs. for 366.2 for not including ethnicity, Wilcox. p = 8.5E-5). However, 10 of 73 participants appeared better fit by a model including POC bias (Fig. 2B, C).

In pre-registration, we hypothesized that across-participant coalitional threat, operationalized as perceiving oneself as of lower socioeconomic rank, would increase harm- intent attributions – but we found evidence for the opposite. Average harm attributions increased with Subjective SES (SSES) score, beta=0.109, Std.Err.=0.048, p=0.030. This was unchanged controlling for testing wave and drug (HIAv ∼ SSES + wave*drug + (1|participant): p=0.034, beta=0.109, Std.Err.=0.050).

### The success of the winning model replicated between baseline and follow-up waves

A winning model over the baseline sample motivated the hypothesis that the same model would be the best at follow-up. This hypothesis held, in that the same model had the best total BIC in the follow-up testing. Model comparison at baseline vs. follow-up is shown at Fig. 4.

We then examined the hypothesis that each parameter of the model would show stability, i.e. that it would be correlated between baseline and follow-up. To do this, we regressed the follow-up values of the parameters on the baseline ones, controlling for group allocation (placebo vs. SSRI). We found evidence for stability of the following parameters: *initHarmInt* (p=0.0096, adj.R^2^ = 0.096 excl. one outlier; see Methods), decision noise *decisPrec* (p=0.0024, adj.R^2^ =0.11), prior attribution certainty *typingConf* (p=0.00011, adj. R^2^ = 0.21 excl. one outlier), bias *fairnessB* (p=1.5E-5, adj. R^2^ = 0.253 excl. one outlier; Note this is *not* bias about ethnicity but propensity to attribute beneficence), memory *learnRet* (p=0.00122, R^2^ =0.132), and learning over dictators *λ*_other_ (p=0.012, R^2^ =0.071). Other parameters were poorly correlated (pS0: p=0.194, R^2^ =0.064; dEv: p=0.731, adj.R^2^ = -0.032 excl. one outlier; wH: p=0.806, R^2^ =0.010), wS (p=0.436, R^2^ =-0.022). As often found, model fit was the most stable measure (Log-likelihood: p=2.17E-6, adj. R^2^ = 0.287) and improved on re-testing (See Fig. S6, B. vs. A.)

### Including predictions along with attributions improves task and model stability

We then fitted the winning model only to the harmful intent and self-interest attributions (i.e., like the previously published version of the task). In support of pre-registered hypothesis C., fewer measures were correlated at conventional levels of significance, and baseline values generally explained less variance of the follow-up ones. As expected, model fit was most stable (log likelihood: p=0.0074, adj. R^2^ = 0.085) and the overall bias *fairnessB*, decision noise *deciPrec* and learning over dictators *λ*_other_ also showed significant correlations (w0: p=0.012, adj. R^2^ = 0.106; alphaPrec: p=0.00202, adj. R^2^ = 0.122; *λ*_other_: p=0.0208, adj. R^2^ = 0.055; Fig. S5). However, the other measures did not attain conventional significance by this simple measure (p-value for initHarmInt: 0.235, initSelfInt: 0.90, aEv: 0.814, typingConf:0.108, wHarmInt: 0.228, wSelfInt: 0.948, learnRet: 0.155).

### Citalopram may reduce typing confidence, thus enhancing refinement of views

We first assessed the stability of the measure of average attribution levels for an individual per testing wave, as we pre-registered the hypothesis that Citalopram would reduce this measure levels across waves. These measures showed good stability, as shown by correlating follow-up attributions with baseline ones, while controlling for the effect of Citalopram. Average harm- intent attributions, HIAv, correlated at p=0.0015, beta=0.442, R^2^ = 0.135. Average Self-interest, SIAv, showed p= 0.0023, beta=0.409, R^2^ = 0.152. Predictions (predAv) showed p=1.17E-5, beta=0.437, R^2^ = 0.249.

We used linear mixed effects analysis ( measure ∼ wave*drug + ( 1 | participant) ) to assess the effect of Citalopram. We found no evidence that Citalopram affected harm-intent attributions, p=0.11 for the wave*drug term for measure=HIAv. There was modest evidence that self-interest attributions were *enhanced* by Citalopram ( wave*drug for SIAv: beta = 0.269, Std.Err.=0.134, p =0.0484). Average expectation was unaffected, p for predAv = 0.48.

We found a possible novel effect of Citalopram on the confidence of typing characters, typingConf. Otherwise, modeling measures were consistent with the analysis of average attributions, i.e., there was a modest effect of SSRI increasing initSelfInt (pSI0: beta=1.32, Std.Err.=0.62, p=0.037), consistent with SIAv above. The novel effect was a decrease typingConf by Citalopram (aEv: wave*drug p=0.023, beta=-1.18, St.Err.=0.50). This would enhance the attribution-refining process. Notably, this was on a background of initSelfInt reducing with wave (pS0: p=0.014, beta=-2.612, Std.Err.=1.033), unlike initHarmInt (pH0: p=0.153). The wave*drug effect for other measures was unremarkable (p values: LL: 0.729, initHarmInt: 0.217, dEv: 0.743, alphaPrec: 0.7638, fairnessB: 0.568, wHarmInt: 0.979, wS: 0.11, ω: 0.303, λ_other_: 0.898).

Guessing which group participants were in was not associated with initSelfInt (p= 0.24), SIAv (p= 0.60) or typingConf (p= 0.76), hence expectation effects are unlikely to account for our findings.

### Psychometric findings and the effect of Citalopram

We administered the Hypomanic Personality Scale (HPS) at baseline, and the Patient Health Questionnaire (PHQ-9), Rosenberg Self Esteem (RSE), Mood and Anxiety Symptom Questionnaire (MASQ), Perceived Stress Scale (PSS), Generalized Anxiety Disorder - 7 (GAD), and the Achievement Motivation Inventory (AMI) at both baseline and follow-up. We examined the dependence of follow-up scores on Citalopram treatment, controlling for baseline scores, gender, SSES and HPS. Citalopram had no statistically significant effect on any of the scores (Supplemental Fig. S4).

In order to reduce dimensionality and increase sensitivity, we performed a factor analysis on the baseline state-like symptoms (i.e., not the HPS), resulting in two factors on the basis of parallel analysis and scree plot (Fig. S6). We tested the validity and stability of this by deriving factor scores on baseline and follow-up based on the baseline loadings only. The first factor scores, ‘anxious depression’ correlated *r*=0.791, p < 1E-10. The second factor scores, ‘stressful amotivation’, correlated at *r*=0.803, p < 1E-10, both showing good stability. Anxious depression was not affected by SSRI (anxDep ∼ wave*drug + (1 | participant) gave p=0.759 ) but stressful amotivation marginally improved, p=0.052, beta = -0.25. Exploratory linear regressions found no evidence that variance in psychometric ratings, or changes in these ratings, was related to model parameters.

## Discussion

We sought to refine our understanding of the computational basis of attributing motives and examined the role of Serotonin and ethnicity in such attributions. We tested whether SSRI treatment may promote attributions of more beneficent motives to others; and whether factors often postulated to recruit subtle coalitional dynamics, namely apparent race and subjective socioeconomic status, affected attribution of motives. Importantly, we found strong evidence in favor of our hypothesized ’Classify-refine’ models, which postulated that participants rapidly classified others’ attributes as positive or negative, and then sought to refine attributions.One week’s treatment with Citalopram did not result in more magnanimous attributions (Moutoussis et al., 2022a), but exploratory evidence suggested that it rendered the process of belief refinement more flexible. Contrary to our hypotheses regarding ethnicity, apparent race did not affect positive or negative attributions, consistent with most of our participants’ decision-making being unbiased in this respect. Notably, low subjective socioeconomic status reduced rather than increased attributions of harmful intent.

In line with our pre-registered modeling hypotheses, we replicated the set of features, or parameters, needed to model the data according to our previous work - with one exception (Barnby et al., 2022). That is, prior belief uncertainty over Harm-intent and Self-interest could be condensed into a single uncertainty parameter. This is important, as the replication entails a task in a different population, a laboratory based pharmacology experiment, with added task features of naturalistic photos of the ’partner’, and questions about expected frequency of fair outcomes (Fig. 1). We found good evidence that the version of the task including these new questions was more stable, or reliable, than the original - as we predicted ((Moutoussis et al., 2022a), main hypothesis C). Our close replication of a robust correlation between initial beliefs in harm-intent and self-interest (Barnby et al., 2022) suggests two things. First, that there is a general propensity to attribute beneficent vs. detrimental motives. Second, that future studies should consider their commonality (harm-benefit intent) and contrast (other-self focus). Here we used active-inference, which offers a natural framework for our models, but our modeling findings can be implemented in other frameworks (like reinforcement learning) too.

The winning Classify-refine learning shares much with posterior-belief-based credit allocation models (Dorfman et al., 2019), and has been considered in some detail by Story and co-authors in the context of ’black and white thinking’ (Story et al., 2023). Refining this thinking, we suggest that classify-refine updating may be a useful socio-cognitive strategy, rather than a testing artifact, or mostly found in suboptimal black-and-white thinking. It is advantageous to rapidly distinguish friend from foe, and then do more justice to the true nature of others. It may thus be an example of benign "thinking fast and slow" (Kahneman, 2011).Classify-refine may also explain the fast-then-slow social learning shown by Bone, Pike and co-workers, and others (Bone et al., 2021; Hopkins et al., 2021). Bone et al fitted their data with associative models wherein learning rates decayed in novel ways, partially inspiring the present work. Future research should compare our (Bayesian) classify-refine models with associative learning employing multiple learning processes, by which the brain may approximate Bayesian inference. But how might classify-refine relate to the well recognised, maladaptive polarized thinking? This could be about the failure of the ’refine’ process, either due to excessive ’typing confidence’ leading to persistent stereotyping, or by circular causation induced by tit-for-tat strategies.

Citalopram treatment did not increase attribution of more prosocial motives, either in terms of increasing the proportion of fair returns that participants expected, their average HI and SI attributions, nor the parameters pHI0 and pSI0. Indeed, there was weak evidence for a relative increase in average SI and pSI0 (both at the 0.05 level uncorrected). Citalopram did not affect our healthy participants’ psychometric measures either, so the possibility remains that it may only ameliorate truly depressive attributions, and/or over a longer timecourse. We found intriguing exploratory evidence – much in need of replication – that Citalopram may reduce certainty over character types, allowing for faster learning of others’ true nature, and that it may reduce stress-induced amotivation. The simplest interpretation of our findings, however, is that treatments for depression that ameliorate inter-personal cognition may act via pathways only peripherally, or indirectly, affected by antidepressants (Nord et al., 2021).

Important calls have been made in recent years towards a translational neuroscience sensitive to its own biases, one which can elucidate and help mitigate prejudice (Iyer, 2022; Kaiser Trujillo et al., 2022; Singh et al., 2022). Our findings about the role of race were unexpected, and in a sense cause for celebration. Models that included a parameter that would change motivational attributions depending on the ethnicity of the partner, a parameter which was able to capture any direction of such effect at the individual level, were rejected as they did not improve model fit – although a few individuals may be prone to this bias, and they may have a disproportionate impact on disadvantaged individuals. Our findings contrasts with literature showing affective biases against people of color (Davies & Turnbull, 2011) and is consistent with decision-making findings where appropriately educated participants behaved fairly towards POC others, despite having low-level affective biases (Correll et al., 2007). Lower socio-economic self-ranking was associated, again contrary to our hypotheses, with lower attributions of harm intent, a positive finding in the sense of lack of evidence for excessive mistrust in this group. We note that our participants were predominantly young, highly educated women from BAME backgrounds, the most common ethnicity being Chinese. Groups with a different experience of social disadvantage, or indeed advantage, may attribute motives differently. Although, therefore, our sample is not representative of the UK population, our inability to detect signs of attribution bias and socioeconomic-based suspiciousness are cause for measured celebration.

In terms of limitations, our study did not involve participants with clinically significant symptoms, nor did Citalopram have an effect on their mood and anxiety, rendering moot the question of whether the attributional problems of depression are ameliorated by SSRIs. It would be important, therefore, for CBT treatment trials to include social-cognitive tasks such as the one used here. Similarly, the happy absence of detectable biases related to low subjective socio- economic status or ethnicity need replication in samples more representative of the general population.

In conclusion, Classify-refine models appear to hold much promise for social neuroscience, have normalizing implications for ’black-and-white’ thinking, are well served by the active inference framework, and have learning-relevant parameters that may depend on Serotonin function. SSRIs appear to have little effect on the motives and expectations healthy people have from each other. Importantly, computational neuroscience studies should be equipped to interrogate burning issues such as social inequality or racial biases, but, as importantly, also to elucidate mechanisms behind ‘the glass being half full’.

## Supporting information

Supplement with method details

## Acknowledgments

We wish to thank Giles Story, Alex Pike, Kate Button, Janina Hoffman, Katie Hobbs, Ryan Smith and the WCHN Methods group for their support.

## Funding

The Department of Imaging Neuroscience is supported by a Platform Grant from the Wellcome Trust, and its Max-Planck center for computational neuroscience and ageing is supported by UCL and the Max-Plan Society. R.B.R. is supported by a NARSAD Young Investigator Award from the Brain & Behavior Research Foundation, P&S Fund. L.M. is supported by a Medical Research Council Clinician Scientist Fellowship (MR/S006613/1).

## Competing Interests

MC is a current employee of and holds options in COMPASS Pathfinder Ltd., but this work is unrelated to COMPASS Pathfinder. None of the other authors has any financial or intellectual competing interests in connection with the manuscript to declare.

## Author Contributions

**Michael Moutoussis:** investigation, data curation (supporting), review & editing (equal), conceptualization, formal analysis, methodology, software, writing – original draft preparation (all lead), **Joe Barnby:** Conceptualization, formal analysis, methodology, software, writing – original draft preparation (all supporting), review & editing (equal). **Anais Durand:** Investigation, data curation (equal), writing – original draft preparation (supporting), review & editing (equal)**, Megan Croal** Investigation, data curation (equal), writing – original draft preparation (supporting), review & editing (equal)**, Laura Dilley:** Supervision, writing (supporting), review & editing (equal), **Robb B. Rutledge:** Methodology, project administration, supervision, writing – original draft preparation (all supporting) review & editing, (equal), funding acquisition (lead). **Liam Mason** Conceptualization, methodology, writing – original draft preparation, investigation, data curation, funding acquisition (all supporting), review & editing (equal), project administration, supervision (both lead).

## Notes

https://github.com/mmoutou/Classify-refine_Sharing_game

### Summary of Updates

Substantial revisions were made after the first round of peer reviews upons submission for publication.

## References

1. Barnby, J. M., Bell, V., Mehta, M. A., & Moutoussis, M. (2020). Reduction in social learning and increased policy uncertainty about harmful intent is associated with pre-existing paranoid beliefs: Evidence from modelling a modified serial dictator game. PLOS Computational Biology, 16(10), e1008372. 10.1371/journal.pcbi.1008372

2. Barnby, J. M., Mehta, M., & Moutoussis, M. (2022). The computational relationship between reinforcement learning and social inference in paranoia. PsyArXiv. 10.31234/osf.io/x4d3f

3. Bone, J., Pike, A. C., Lewis, G., Lewis, G., Blakemore, S.-J., & Roiser, J. (2021). Computational mechanisms underlying social evaluation learning and associations with depressive symptoms during adolescence. PsyArXiv. 10.31234/osf.io/9m7vr

4. Brown, G. D. A., Lewandowsky, S., & Huang, Z. (2022). Social sampling and expressed attitudes: Authenticity preference and social extremeness aversion lead to social norm effects and polarization. Psychological Review, 129(1), 18–48. 10.1037/rev0000342

5. Correll, J., Park, B., Judd, C. M., Wittenbrink, B., Sadler, M. S., & Keesee, T. (2007). Across the thin blue line: Police officers and racial bias in the decision to shoot. Journal of Personality and Social Psychology, 92, 1006–1023. 10.1037/0022-3514.92.6.1006

6. Davies, J. L., & Turnbull, O. H. (2011). Affective bias in complex decision making: Modulating sensitivity to aversive feedback. Motivation and Emotion, 35(2), 235–248. 10.1007/s11031-011-9217-x

7. Dorfman, H. M., Bhui, R., Hughes, B. L., & Gershman, S. J. (2019). Causal Inference About Good and Bad Outcomes. Psychological Science, 30(4), 516–525. 10.1177/0956797619828724

8. Greenburgh, A., Bell, V., & Raihani, N. (2019). Paranoia and conspiracy: Group cohesion increases harmful intent attribution in the Trust Game. PeerJ, 7, e7403. 10.7717/peerj.7403

9. Hopkins, A. K., Dolan, R., Button, K. S., & Moutoussis, M. (2021). A Reduced Self-Positive Belief Underpins Greater Sensitivity to Negative Evaluation in Socially Anxious Individuals. Computational Psychiatry, 5(1), Article 1. 10.5334/cpsy.57

10. Iyer, C. (2022). ‘Neuralizing’ Injustice: How neuroscience misunderstands racism, addiction, and crime. *Intersect: The Stanford Journal of Science*, Technology, and Society, 16(1), Article 1. https://ojs.stanford.edu/ojs/index.php/intersect/article/view/2250

11. Joseph, N. T., Myers, H. F., Schettino, J. R., Olmos, N. T., Bingham-Mira, C., Lesser, I. M., & Poland, R. E. (2011). Support and Undermining in Interpersonal Relationships Are Associated with Symptom Improvement in a Trial of Antidepressant Medication. Psychiatry: Interpersonal and Biological Processes, 74(3), 240–254. 10.1521/psyc.2011.74.3.240

12. Kahneman, D. (2011). *Thinking, Fast and Slow*. Farrar, Straus and Giroux. https://books.google.co.uk/books?id=ZuKTvERuPG8C

13. Kaiser Trujillo, A., Kessé, E. N., Rollins, O., Della Sala, S., & Cubelli, R. (2022). A discussion on the notion of race in cognitive neuroscience research. Cortex, 150, 153–164. 10.1016/j.cortex.2021.11.007

14. Maina, I. W., Belton, T. D., Ginzberg, S., Singh, A., & Johnson, T. J. (2018). A decade of studying implicit racial/ethnic bias in healthcare providers using the implicit association test. Social Science & Medicine, 199, 219–229. 10.1016/j.socscimed.2017.05.009 matlab. (2019). *MATLAB* (9.7) [Computer software]. The MathWorks Inc. https://www.mathworks.com

15. Moutoussis, M., Barnby, J. M., Durant, A., Croal, M., Rutledge, R., & Mason, L. (2022b). Do SSRIs promote more magnanimous attributions about others? 10.17605/OSF.IO/HT95F

16. Moutoussis, M., Barnby, J. M., Durant, A., Croal, M., Rutledge, R., & Mason, L. (2022a). Do SSRIs promote more magnanimous attributions about others? 10.17605/OSF.IO/HT95F

17. Moutoussis, M., Hopkins, A. K., & Dolan, R. J. (2018). Hypotheses About the Relationship of Cognition With Psychopathology Should be Tested by Embedding Them Into Empirical Priors. Frontiers in Psychology, 9. 10.3389/fpsyg.2018.02504

18. Nord, C. L., Barrett, L. F., Lindquist, K. A., Ma, Y., Marwood, L., Satpute, A. B., & Dalgleish, T. (2021). Neural effects of antidepressant medication and psychological treatments: A quantitative synthesis across three meta-analyses. The British Journal of Psychiatry, 219(4), 546–550. 10.1192/bjp.2021.16

19. R Core Team. (2020). R: A language and environment for statistical computing. R Foundation for Statistical Computing, Vienna, Austria*. URL* http://www.R-project.org/. [Computer software].

20. Raihani, N. J., & Bell, V. (2017). Paranoia and the social representation of others: A large-scale game theory approach. Scientific Reports, 7(1), Article 1. 10.1038/s41598-017-04805-3

21. She, Z., Ng, K.-M., Hou, X., & Xi, J. (2022). COVID-19 threat and xenophobia: A moderated mediation model of empathic responding and negative emotions. Journal of Social Issues, 78(1), 209–226. 10.1111/josi.12500

22. Singh, M. K., Nimarko, A., Bruno, J., Anand, K. J. S., & Singh, S. P. (2022). Can Translational Social Neuroscience Research Offer Insights to Mitigate Structural Racism in the United States? Biological Psychiatry: Cognitive Neuroscience and Neuroimaging, 7(12), 1258–1267. 10.1016/j.bpsc.2022.05.005

23. Smith, H. L., Summers, B. J., Dillon, K. H., Macatee, R. J., & Cougle, J. R. (2016). Hostile interpretation bias in depression. Journal of Affective Disorders, 203, 9–13. 10.1016/j.jad.2016.05.070

24. Smith, R., Friston, K. J., & Whyte, C. J. (2022). A step-by-step tutorial on active inference and its application to empirical data. Journal of Mathematical Psychology, 107, 102632. 10.1016/j.jmp.2021.102632

25. SPM development team. (2022). *SPM* (Version 12) [Matlab/octave]. https://www.fil.ion.ucl.ac.uk/spm/software/spm12/

26. Stjohn, C., & Healdmoore, T. (1995). Fear of Black Strangers. Social Science Research, 24(3), 262–280. 10.1006/ssre.1995.1010

27. Story, G. W., Smith, R., Moutoussis, M., Berwian, I. M., Nolte, T., Bilek, E., Siegel, J. Z., & Dolan, R. J. (2023). A social inference model of idealization and devaluation. Psychological Review, No Pagination Specified-No Pagination Specified. 10.1037/rev0000430

