## Supplement with method details for "Rapid classification vs. refined learning about harm intent: Roles of serotonin and racial bias"

**This Supplement includes:**

Supporting text: 'A. Supplementary Methods'

Figures S1 to S7

Tables S1, S2 and S3

SI References

**Other supporting materials for this manuscript will be made available for reviewers:**

Dataset S1

Software S1

### A. Supplementary Methods

Intuition to the workings of the classify-refine attribution model are given in Fig. S1, then a detailed active-inference formulation of models used here follows. We note that the 'classic' active inference models used here are as precise an adaptation as practicable of our previous work (Barnby et al., 2022) into the language of active inference.

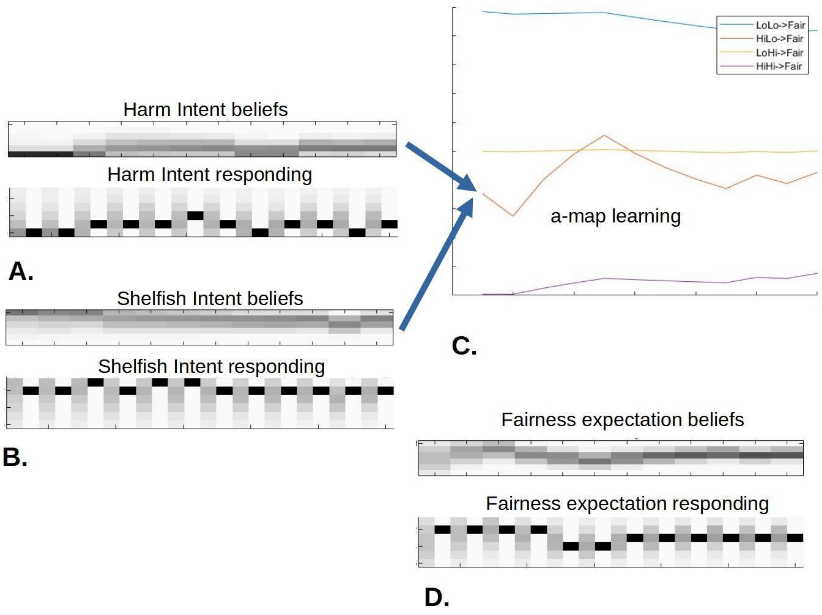


**Figure S1.** Demonstration of the workings of the classify-refine model. All x-axes are 12 trials. Here, the agent starts with a strong prior belief that the Dictator has very low harm intent, but high self-interest. This is why the bottom panels of **A.** and **B.** start from a dark band at low and high values respectively. A. and B. in fact show the overall belief of the average of each attribute. This is translated by the class-based beliefs by considering both the beliefs about high or low attributes, and the amount of effective evidence that supports them. Little evidence results in less certain beliefs. The bottom panels of A. and B show the initial beliefs about declaring a particular level of the attribute, and the posterior about this action (gray -> dark in each trial). **C.** shows the likelihood the agent holds for each combination of attributes (High SI + high HI, high-SI low-HI, etc.) to result in a fair split. Because the agent is confident that the high-SI low-HI disposition obtains, they mostly modify their beliefs about the Hi-Lo likelihood. **D.** The agent's resulting beliefs, and reports, that a fair split will obtain. Note that the fairness beliefs become more and more confident with trials (darker, steady band) while each attribute belief has become less certain.

#### The likelihood of fair and unfair splits under different levels of Harm-intent and Self-interest

The learning core of the model was formulated as a Hidden Markov process, which did not infer actions. The process model (see main Fig 2A) contained two state factors, {1} = Harm-intent, {2}

= Self-interest.

In the classical models, as in our previous work, states for each factor came from a relatively fine-grained scale from low to high, containing Ng bins for each dimension of attribution. Thus the state spaces for HI and SI were grids *g* = {1/(2*Ng*), 3/(2*Ng*) ... (2*Ng*-1)/(2*Ng*)} . After preliminary exploration, we chose *Ng*=6. In all models, participants held belief mixtures, not all-or-nothing beliefs, over such attribute states of the Other.

Participants mapped levels *HIj* , *SIk* to the resulting probability of fair outcome through a logistic sigmoid, containing weights *wH*, *wS*, *w0,* which were fixed parameters for each participant (Eq. 1).

$$\pi_{fair}(HI_j, SI_k) = \texttt{invlogit}((\frac{1}{2} - HI_j)\, wH + (\frac{1}{2} - SI_k)\, wS\; - w0) \qquad \text{Equation S1}$$

Still in the classic models and in active-inference notation, Equation S1 was used to populate a generative likelihood map *A,* which assigned a probability of fair outcome for each possible combination of HI, SI.

In classify-refine models, the central core considered only two states along each attribution dimension, giving four combinations, *s*Low-Low , *s*Low-High etc.. Initially, these corresponded to Low = 0.05 and High = 0.95 levels, so as to represent the psychological meaning of low and high levels of each attribute, and to span a most of the range of the attributes. However, the likelihood map from states to outcomes, here denoted as *a,* did not contain probabilities directly. It contained concentration parameters, or amounts of effective evidence for the relevant (beta belief) distributions over outcomes. Their initial values, representing the confidence or precision that a state of particular label, say *s*Low-High would lead to a fair outcome was proportional to the *typing confidence* parameter *aEv* . This is shown in Table S1.

| **Table S1** | | | | | |
|---|---|---|---|---|---|
| State label | State factor 1 level | State factor 2 level | Initial values of HI, SI | Initial concentration parameters (beta parameters) of likelihood map | |
| | Harm intent | Self interest | | **Fair outcome** | **Unfair outcome** |
| *s*11 | Low | Low | **0.05 , 0.05** | aEv . *π*fair ( *s*11 ) | aEv . (1-*π*fair ( *s*11 ) ) |
| *s*12 | Low | High | **0.05 , 0.95** | aEv . *π*fair ( *s*12 ) | aEv . (1-*π*fair ( *s*12 ) ) |
| *s*21 | High | Low | **0.95, 0.05** | aEv . *π*fair ( *s*21 ) | aEv . (1-*π*fair ( *s*21 ) ) |
| *s*22 | High | High | **0.95, 0.95** | aEv . *π*fair ( *s*22 ) | aEv . (1-*π*fair ( *s*22 ) ) |

To put *A* (or *a*) into practice, each trial included two time-steps, namely initial state and split-observation-state.

**Refining the likelihood map through learning**

In the classify-refine model, after each observation the participant can refine their likelihood map. To develop intuition to this, imagine a participant who uses Eq. 1 to estimate that initially

$\pi$fair ( $s$22={0.95,0.95} ) = 0.05 . They then observe a long series of outcomes, 0.15 of which are fair. This may still be consistent of the Dictator being in $s$22 , i.e. of high HI and SI, but the likelihood function should now predict ~ 0.15 fair outcomes. In our active inference framework, to update the likelihood map is easy - the concentration parameters for each cell of the map are augmented by their estimated responsibility in causing said outcome.

If we denote by $st$ the vector of beliefs at the end of trial $t$ that each state may have obtained, the accumulation of the a - map entries is written succinctly in terms of an outer vector-product. We also include a memory parameter ω to account for forgetting between trials:

$$\boldsymbol{a}_t = \omega\, \boldsymbol{a}_{t-1} + o_t \otimes s_t \quad \text{Equation S2}$$

This is standard active inference, but here, once the likelihood map is updated, what are the new values of HI and SI consistent with it? To do this, we minimized a distance measure between { $HI$low , $SI$low , $HI$high , $SI$high } and the updated a-map as follows:
a. Tabulate a larger grid $G$ of all possible values of { $HI$low , $SI$low , $HI$high , $SI$high } allowed by component grids identical to the above $g$ ={1/12, 3/12 ... 11/12} .
b. Calculate the resulting policy for each cell of $G$ using Eq. 1

c. Calculate the squared differences $\Delta$ between the fair outcome probabilities in each cell of the updated a-map and the corresponding ones in $G$.

d. Weigh each element of $\Delta$ by the evidence, i.e. the sum of the relevant concentration parameters in the a-map, on which it was based.

e. Select the element of G nearest to the a-map according to this weighed sum-square-difference.

*The reporting mechanisms*

Each of the attributional reporting processes (Fig. 2C) formed a distribution over the $Ng$ report bins. We estimated the expected HI as follows:

$$HI_m = \sum_{i,k \in \{low, high\}} \frac{d(s_{ik})}{d_{total}} HI_i \quad \text{Equation S3}$$

Where $d$($s$) and $dtotal$ are counts in evidence of the state $s$ and for all the states. We then approximated discrete beliefs over the $Ng$ bins using a cumulative beta density :

$$\begin{aligned} d_{HI}(1) &= CDF(\,Beta(g(1); HI_m\, d_{total}, (1 - HI_m)\, d_{total})) \\ for\ 1 < j &\leq Ng\ : \\ d_{HI}(j) &= CDF(\,Beta(g(j); HI_m\, d_{total}, (1 - HI_m)\, d_{total})) - \sum_{k<j} p_{HI}(k) \end{aligned}$$

Equation S4

The distribution over SI is computed similarly.

Reporting beliefs about the probability of fair return was similar, but instead of Eq. 4, the expectation of fair return was formed out of the a-map concentration parameters:

$$\begin{aligned} d_{fairness}(1) &= \sum_{o=fair,\ all\ \ j,k} a_{o,j,k} \\ d_{fairness}(2) &= \sum_{o=unfair,\ all\ \ j,k} a_{o,j,k} \end{aligned}$$

Equation S5

Reporting was then carried out by a two-step active-inference MDP, where report-states, and hence the actions leading to them, were more desirable the closer they were to the most probable (highest *d* in Eq. 4,5) belief. We note that as per Fig. 2, the 'response spokes' received, but did not send, information to the learning 'core'. This means that the overall model should not be seen as a single active-inference MDP, and that free energy was optimized in the 'response' and 'core' processes separately.

**Learning and coalitional dynamics from partner to partner**

The parameters *initHarmInt* and *initSelfInt* (*pHI0 and pSI0*), which controlled prior beliefs about Dictators, were modified to model learning and coalitional dynamics between partners. To affect learning, the shift in probability of Harm Intent, *pHI* (and *pSI* ) from prior to posterior was calculated, and a fraction *λother* of it used to update *pHI0, pSI0* :

$$\begin{aligned} \delta pHI &= \vec{d}_{HI} \cdot \vec{g} - pHI0 \\ (\texttt{ditto} \quad & \texttt{for} \quad \delta pSI) \\ pHI0 &\leftarrow pHI0 + \lambda_{other}\, \delta pHI \\ dEv &\leftarrow dEv + \lambda_{other} \\ aEv &\leftarrow aEv + \lambda_{other} \end{aligned}$$

Equation S6

In contrast, the coalitional dynamics did not affect partner-to-partner learning, but simply modulated the pHI0 and pSI0 for the partner in question as per

$$\begin{aligned} pHI0' &\leftarrow pHI0^{POCbias} \\ pSI0' &\leftarrow pSI0^{POCbias} \end{aligned}$$

Equation S7

This allows modeling of the coalitional bias of each individual in whichever direction this may be - including having making more optimistic attributions for their out-group.

**The transition map, or B-matrix, utility map, *C*, and prior beliefs over states, *d***

A transition map B (in active inference notation) with *B*{1} = *I* and *B*{2} = *I* . *I* is the identity matrix mapping, mapping each of the probabilities of the states in each factor, to the same values at the next step. For Classify-refine,

$$B\{1\} = B\{2\} = I(2)$$

Equation S8

as there are 2 states per factor.

A utility map *C* giving the preferences of the agent for unfair and fair splits, in terms of desired outcome probability. This scales the desirability of these outcomes against epistemic value (accurate predictions) of the generative model.

Probability vectors *d*{1} and *d*{2} describing the prior distributions over *s*Low-Low , *s*Low-High etc. - but now simply factorized over HI and SI, so that p( *s*Low-Low ) = p( *s*HI Low ) p( *s*SI Low ) , etc. All models, both Classify-refine and classical, learn what the likely state generating observations were by accumulating evidence in d.

Crucially, the initial values of *d* are parameterized as:

$$d\{1\}(1) = pHI0 \,.\, dEv \,.\, EvRat$$

$$d\{1\}(2) = (1\text{-}pHI0) \,.\, dEv \,.\, EvRat$$

$$d\{2\}(1) = pSI0 \,.\, dEv$$

$d\{2\}(2) = (1\text{-}pSI0)\ .\ dEv$ Equation S9

**'Reporting Spoke' processes**

Reporting processes are modelled with a full MDP over two time steps. There is an initial timestep where a Likert-like reporting policy is chosen, here one of six levels of fairness belief, HI or SI; and a final time-stem where the (probabilistic) action is implemented.

Two state factors exist in each spoke, that is, the true state of the world, and the state of the Likert scale. There are two corresponding factors in each possible policy, but the first one is trivial as the agent cannot change the state of the world. These are stored in an array:

$V = V(\text{time-of-action, Likert-action, world-action})$ Equation S10

The B map again has two factors. *B*{1} is identity, as above. *B*{2} takes the MDP from any state and transitions it with very high probability to the corresponding Likert state. E.g. Likert-action 3 brings any initial Likert-state to Likert-state-3, etc.

The *C* map is interesting, in that the desirability it encodes is about landing in a state corresponding to a 'correct' report, give one's beliefs. We therefore considered three outcomes: 3: 'Quite-correct', 2: 'Almost (one step away from quite-) correct', and 1: 'wrong'. An 'indifferent' outcome 4 obtained outside the time of meaningful action. The entries of *C*, i.e. the numerical reward values for these were fixed at [R1 ... R4 ] = [-4, -2, 3, -1].

The first component of *A, A*{1} simply reported where the agent was in its response state - if it was in Likert state 1, it almost certainly observed state 1, etc.

Given these outcomes, *A*[2} described the likelihood of each combination of true-world-state and Likert-report-state to give outcomes of 1 to 4 above. Likert-states of the same value as the true-world-state mapped almost certainly to quite-correct, adjacent ones to almost-correct, etc. This formulation of *A*[2} naturally allows for an additional layer of social-desirability responding, e.g. observing that one has made a 'quite-correct' response if one systematically offers a somewhat lower HI report than one's true beliefs. However it was not necessary to implement such biases here, as they could be subsumed into the parameters of Equation 1 in the main text.

The prior belief vectors $dHI$, $dSI$ were crucial here, as the reporting action took place on their basis. They were estimated via the equation above, that is, set on the basis of the core beliefs just before the report in question. The reporting MDPs therefore did not communicate between themselves from trial to trial, but only via the core.

### B. Additional Results

#### B1. Demographics

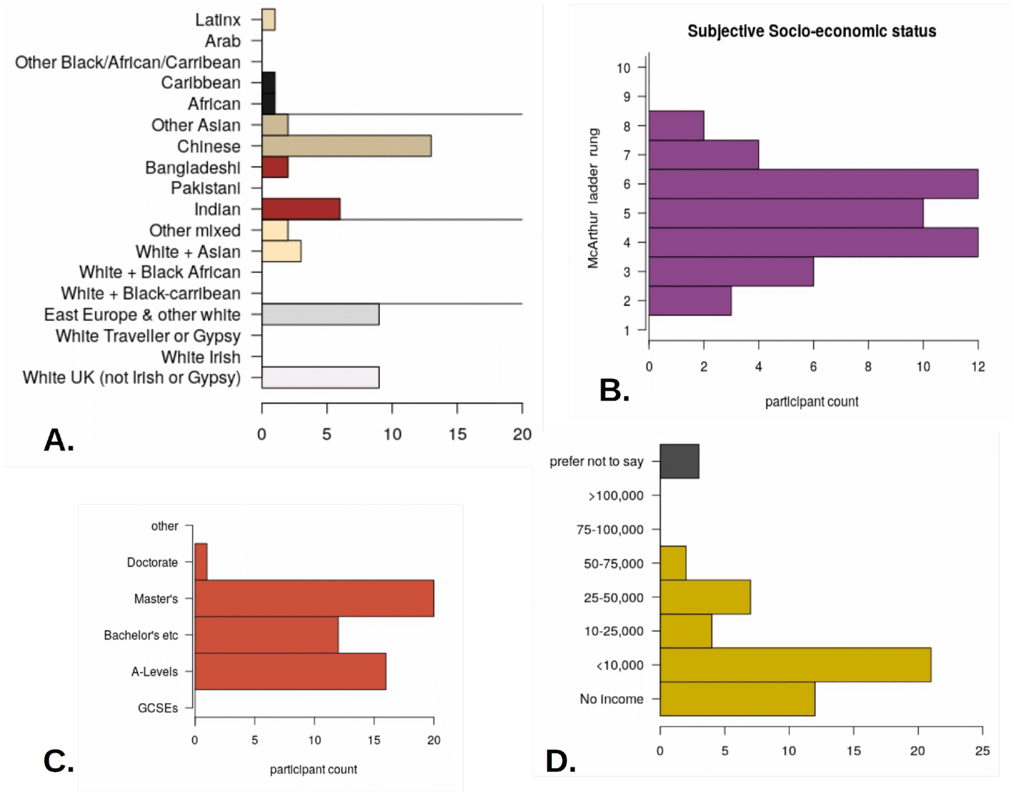


**Figure S2** Demographic characteristics. **A.** About half the sample was between 18 and 25 years old **B.** They perceived their overall socioeconomic status to be around the middle of the population, and this was associated with **C.** high levels of educational achievement, but **D.** very low monetary incomes.

#### B2. Controlling for unblinding

Table S2. Participants significantly guessed whether they were in the Placebo or SSRI groups, but neither this nor their actual group allocation had any effect on drop-out.

| | were given Placebo | were given SSRI |
|---|---|---|
| Guessed Placebo | 18 (58%) | 9 (21%) |
| Guessed SSRI | 4 (13%) | 24 (57%) |
| Could not guess at all (declined, or answered very unclearly) | 9 (29%) | 9 (21%) |
| Pearson Chi-squared: | df=2, X-squared = 16.0, p-value = 0.00034 | |
| Total | 31 | 42 |
| total attending follow-up | 28 | 38 |
| Fisher exact test on drop-outs | Odds ratio = 0.98, p-value = 1 | |

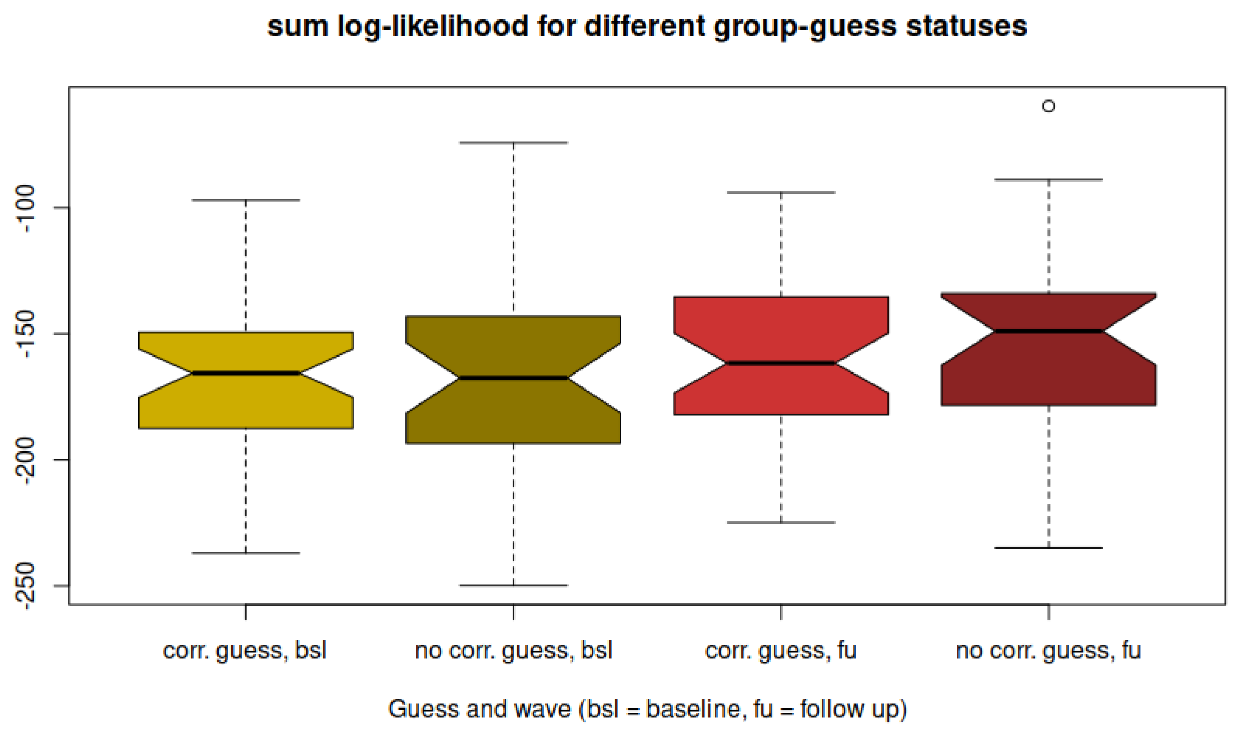


**Figure S3** Neither drug vs. placebo group allocation, not guessing which group the participant was in, was statistically associated with how well the winning model accounted for the data. Notches are centered at the median. Overlapping notches indicate no statistically significant difference.

**B4. Further modeling results**

**Table S3** - Model comparison according to AIC is qualitatively very similar to the one according to BIC, i.e. different form of complexity penalty makes very little difference. Overall winning model highlighted in green.

| **Model** | **Median AIC at baseline** | **Median AIC at follow-up** |
|---|---|---|
| **Models fitted to predictions and attributions** | | |
| Reference 11 parameter. model plus POC bias (model **j**) | 358.6 | 340.5 |
| Reference 11 parameter model without POC bias (**model k**) | 356.7 | 327.0 |
| As per (k) above, but without memory parameter (10 param model, ***l***) | 360.6 | 346.9 |
| As per (k) above, but with wS=wH(Only 1 attribution sensitivity parameter; 10 freely fitted param.; ***m***) | 374.7 | 350.8 |
| As per (k) above, but with wS fixed to its median value (10 freely fitted params, 1 group based param; ***o***) | 354.7 | 340.1 |
| As per (k) above, but no w0 (overall reporting bias; 10 freely fitted param.; ***p***) | 364.8 | 363.1 |
| As per (k), but with equal Self-Interest and Harm-Intent prior uncertainties (10 param, model ***q***) | 354.8 | 325.8 |
| As per (k), but with prior state uncertainty fixed to median of model (k) fit (10 param., model ***r***) | 359.4 | 340.5 |
| As per q, but also with wS fixed to median value (9 freely fitted + 1 group-fitted param; ***s***) | 361.3 | 347.6 |
| **Models fitted to attributions only** | | |
| Same parameterization as winning model ***q*** above | 225.6 | 218.4 |
| As per (u), but wS fixed to its median as per second-best full fit model (9 freely fitted + 1 group fitted param, ***v***) | 221.16 | 272.8 |

### B4. Effects of testing wave and Citalopram

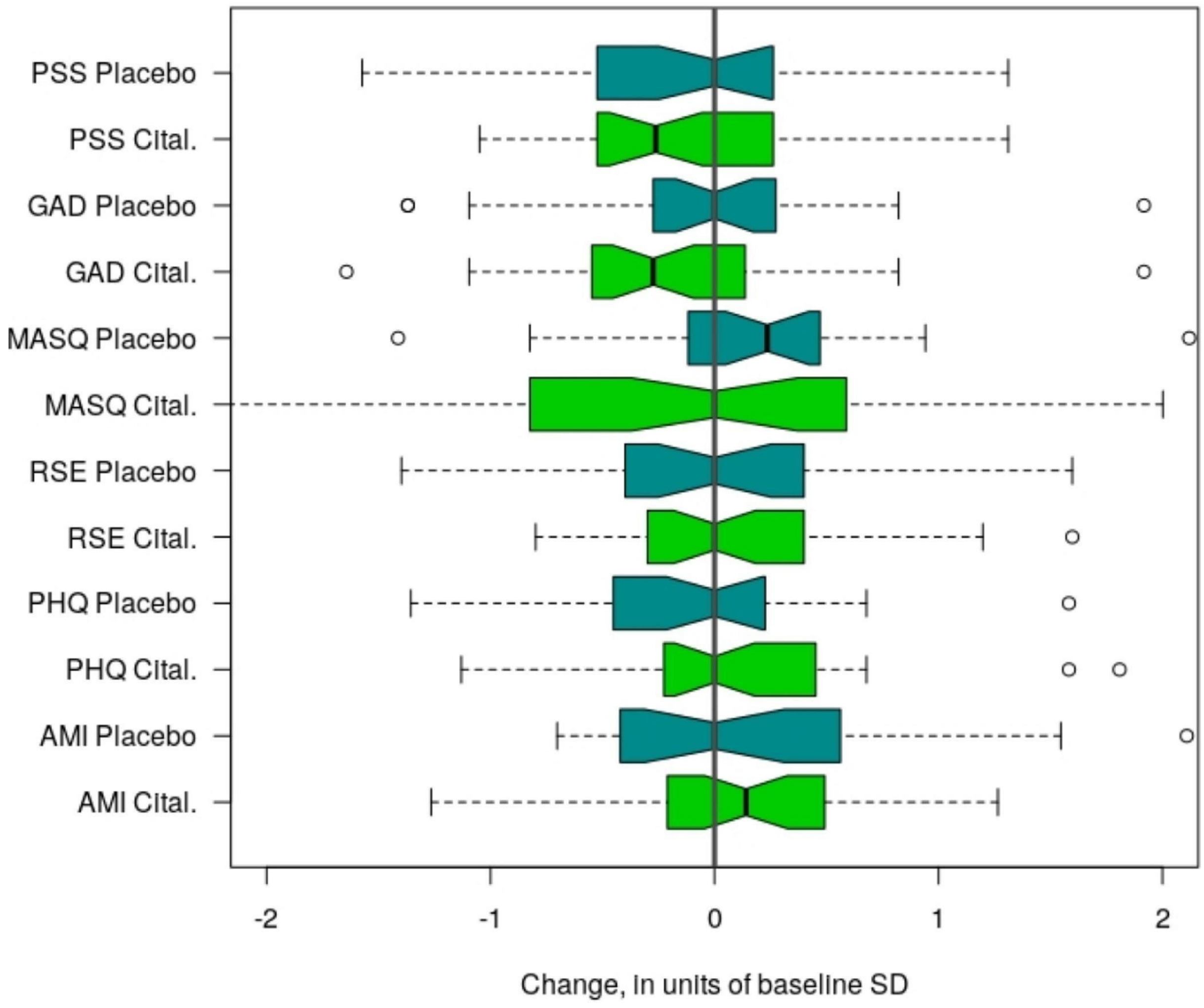


**Figure S4** Raw change from baseline to follow-up for each self-report measure. Darker boxes depict the change for the placebo group, lighter for the Citalopram group. Non-overlapping notches in boxes for the same measure would constitute evidence for significant difference (uncorrected for multiple comparisons) in the medians of the two groups. The notches overlap for all measures. Whiskers extend to +/- 1.5 x interquartile range. P-values for the linear regression coefficients for group described in the main text: AMI: 0.193; PHQ: 0.417; RSE: 0.647; MASQ: 0.347; GAD: 0.905; PSS: 0.361

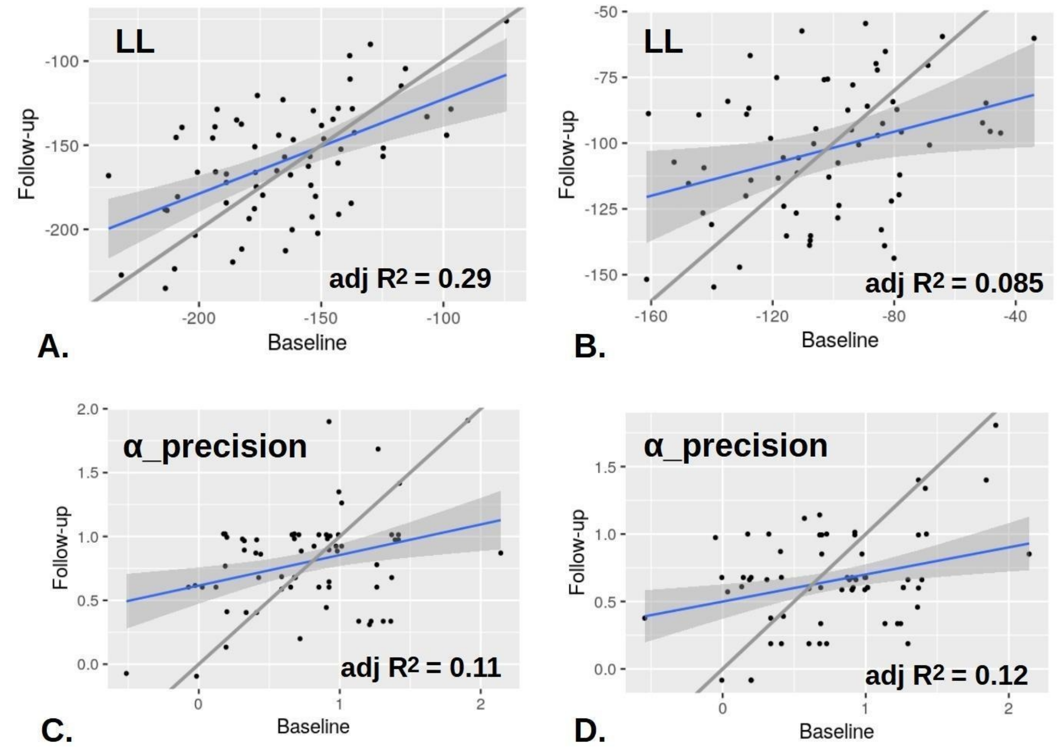


**Figure S5.** Task and model stability. **A., B.** Log-likelihood was the most stable measure, but in the prediction-attribution task (A.) more variance was shared between the baseline and follow-up, typical of the general pattern. However, for the learning over partners and decision noise, here in **C., D.** the attribution only, established task (D.) performed as well.

### B5. Multivariate analyses

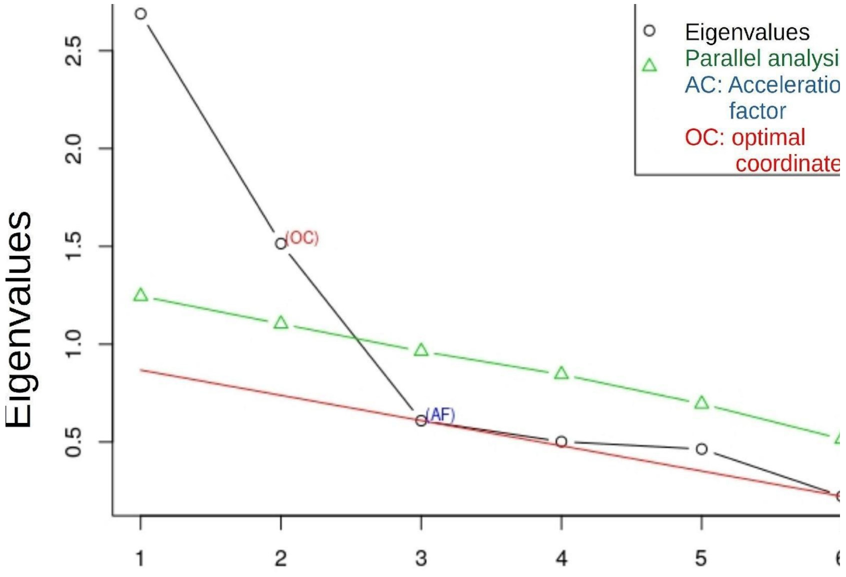


**Figure S6** Screen plot and parallel analysis for psychological symptom factors at baseline.

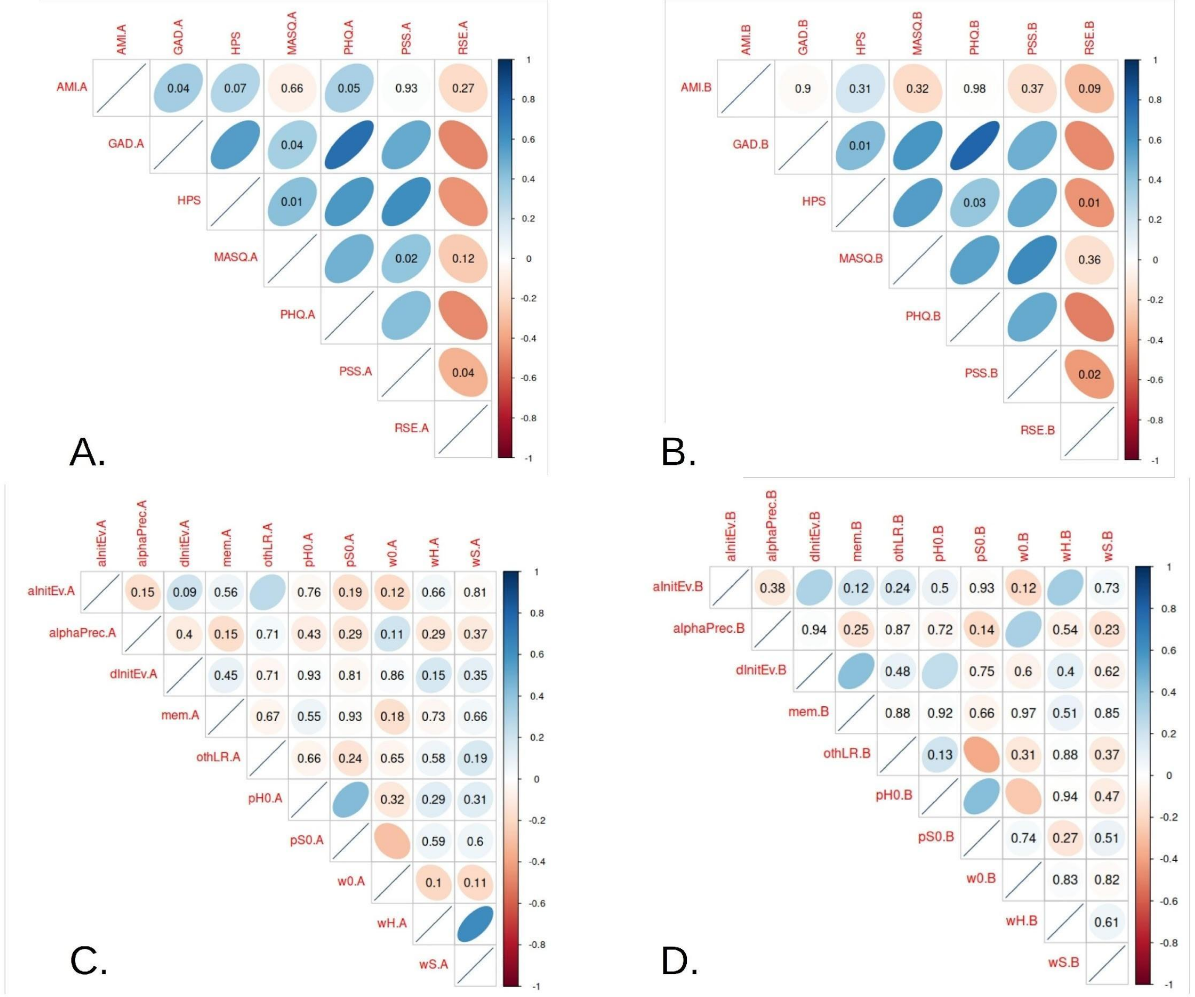


**Figure S7** Correlation matrices for psychological and task-parameter measures. Numbers are uncorrected p-values > 0.05 (non-sig.) **A.** Correlations for the psychological measures in the Citalopram group at baseline (before first dose). Many strong correlations exist, as expected, which form the basis for the factors described in the main text. **B.** The follow-up measures have a very similar structure. C. Correlation matrix for all participants and task parameters at baseline. There are very few significant correlations, even at the uncorrected level. D. Same for follow-up - again, a few significant correlations, and the only one that replicates is the positive correlation between priors on Self-Interest and priors on Harm-intent. This suggests that future work might use beneficence and other-focus, i.e. the mean and difference of Harm-intent and Self-interest.

**Dataset S1 (separate files) and Software S1 (separate files).**

These are available publicly available on github :
https://github.com/mmoutou/Classify-refine_Sharing_game
Instructions to use the functions are found in the README, at
https://github.com/mmoutou/Classify-refine_Sharing_game/blob/main/README.md , the basic ones being:

- To see the basic structure of the model, you may want to step through serialDictator09a.m to see how the basic elements of the MDP (active inference) model are defined A - likelihood map, p(outcome | state) B - Transition map p(next state | current state, action ) C - Goal priors map ( preferences for outcomes - somewhat analogous to rewards in reinforcement learningT ) D - prior probabilities over states

- To see the full likelihood function, you may want to step through spm_mdp_L_vi.m (and versions other than vi). This includes, apart from the basic structure, how learning takes place between blocks and how parameters that relate to ethnicity bias may change the basic parameters.

- To see how the models were fitted using an adaptive grid, see matlab functions like attrssri_Grid09q.m This is the one for the winning model, and is the best commented one.

**SI References**


Barnby, J. M., Bell, V., Deeley, Q., Mehta, M., & Moutoussis, M. (2023). *D2/D3 dopamine supports the precision of mental state inferences and self-relevance of joint social outcomes* (p. 2023.05.02.539031). bioRxiv. https://doi.org/10.1101/2023.05.02.539031

Barnby, J. M., Bell, V., Mehta, M. A., & Moutoussis, M. (2020). Reduction in social learning and increased policy uncertainty about harmful intent is associated with pre-existing paranoid beliefs: Evidence from modelling a modified serial dictator game. *PLOS Computational Biology*, *16*(10), e1008372. https://doi.org/10.1371/journal.pcbi.1008372

Barnby, J. M., Mehta, M., & Moutoussis, M. (2022). *The computational relationship between reinforcement learning and social inference in paranoia*. PsyArXiv. https://doi.org/10.31234/osf.io/x4d3f

Bone, J., Pike, A. C., Lewis, G., Lewis, G., Blakemore, S.-J., & Roiser, J. (2021). *Computational mechanisms underlying social evaluation learning and associations with depressive symptoms during adolescence*. PsyArXiv. https://doi.org/10.31234/osf.io/9m7vr

Brown, G. D. A., Lewandowsky, S., & Huang, Z. (2022). Social sampling and expressed attitudes: Authenticity preference and social extremeness aversion lead to social norm effects and polarization. *Psychological Review*, *129*(1), 18–48. https://doi.org/10.1037/rev0000342

Correll, J., Park, B., Judd, C. M., Wittenbrink, B., Sadler, M. S., & Keesee, T. (2007). Across the thin blue line: Police officers and racial bias in the decision to shoot. *Journal of Personality and Social Psychology*, *92*, 1006–1023. https://doi.org/10.1037/0022-3514.92.6.1006

Davies, J. L., & Turnbull, O. H. (2011). Affective bias in complex decision making: Modulating sensitivity to aversive feedback. *Motivation and Emotion*, *35*(2), 235–248. https://doi.org/10.1007/s11031-

011-9217-x
Dorfman, H. M., Bhui, R., Hughes, B. L., & Gershman, S. J. (2019). Causal Inference About Good and Bad Outcomes. *Psychological Science*, *30*(4), 516–525. https://doi.org/10.1177/0956797619828724
Hopkins, A. K., Dolan, R., Button, K. S., & Moutoussis, M. (2021). A Reduced Self-Positive Belief Underpins Greater Sensitivity to Negative Evaluation in Socially Anxious Individuals. *Computational Psychiatry*, *5*(1), Article 1. https://doi.org/10.5334/cpsy.57
Iyer, C. (2022). 'Neuralizing' Injustice: How neuroscience misunderstands racism, addiction, and crime. *Intersect: The Stanford Journal of Science, Technology, and Society*, *16*(1), Article 1. https://ojs.stanford.edu/ojs/index.php/intersect/article/view/2250
Joseph, N. T., Myers, H. F., Schettino, J. R., Olmos, N. T., Bingham-Mira, C., Lesser, I. M., & Poland, R. E. (2011). Support and Undermining in Interpersonal Relationships Are Associated with Symptom Improvement in a Trial of Antidepressant Medication. *Psychiatry: Interpersonal and Biological Processes*, *74*(3), 240–254. https://doi.org/10.1521/psyc.2011.74.3.240
Kahneman, D. (2011). *Thinking, Fast and Slow*. Farrar, Straus and Giroux. https://books.google.co.uk/books?id=ZuKTvERuPG8C
Kaiser Trujillo, A., Kessé, E. N., Rollins, O., Della Sala, S., & Cubelli, R. (2022). A discussion on the notion of race in cognitive neuroscience research. *Cortex*, *150*, 153–164. https://doi.org/10.1016/j.cortex.2021.11.007
matlab. (2019). *MATLAB* (9.7) [Computer software]. The MathWorks Inc. https://www.mathworks.com
Moutoussis, M., Barnby, J. M., Durant, A., Croal, M., Rutledge, R., & Mason, L. (2022). *Do SSRIs promote more magnanimous attributions about others?* https://doi.org/10.17605/OSF.IO/HT95F
Moutoussis, M., Hopkins, A. K., & Dolan, R. J. (2018). Hypotheses About the Relationship of Cognition With Psychopathology Should be Tested by Embedding Them Into Empirical Priors. *Frontiers in Psychology*, *9*. https://doi.org/10.3389/fpsyg.2018.02504
Nord, C. L., Barrett, L. F., Lindquist, K. A., Ma, Y., Marwood, L., Satpute, A. B., & Dalgleish, T. (2021). Neural effects of antidepressant medication and psychological treatments: A quantitative synthesis across three meta-analyses. *The British Journal of Psychiatry*, *219*(4), 546–550. https://doi.org/10.1192/bjp.2021.16
R Core Team. (2020). *R: A language and environment for statistical computing. R Foundation for Statistical Computing, Vienna, Austria. URL http://www.R-project.org/.* [Computer software].
Singh, M. K., Nimarko, A., Bruno, J., Anand, K. J. S., & Singh, S. P. (2022). Can Translational Social Neuroscience Research Offer Insights to Mitigate Structural Racism in the United States? *Biological Psychiatry: Cognitive Neuroscience and Neuroimaging*, *7*(12), 1258–1267. https://doi.org/10.1016/j.bpsc.2022.05.005
Smith, R., Friston, K. J., & Whyte, C. J. (2022). A step-by-step tutorial on active inference and its application to empirical data. *Journal of Mathematical Psychology*, *107*, 102632. https://doi.org/10.1016/j.jmp.2021.102632
SPM development team. (2022). *SPM* (Version 12) [Matlab/octave]. https://www.fil.ion.ucl.ac.uk/spm/software/spm12/
Stjohn, C., & Healdmoore, T. (1995). Fear of Black Strangers. *Social Science Research*, *24*(3), 262–280. https://doi.org/10.1006/ssre.1995.1010
Story, G. W., Smith, R., Moutoussis, M., Berwian, I. M., Nolte, T., Bilek, E., & Dolan, R. J. (2021). *A Social Inference Model of Idealization and Devaluation*. PsyArXiv. https://doi.org/10.31234/osf.io/yvu2b
